# Mesophyll cells mediate systemic reactive oxygen signaling during wounding or heat stress

**DOI:** 10.1101/2021.02.02.429427

**Authors:** Sara I. Zandalinas, Ron Mittler

**Author notes:** Corresponding author: Ron Mittler. **Author Contributions:** S.I.Z preformed the experiments. S.I.Z. and R.M. designed the experiments, analyzed the results, and wrote the manuscript. All authors edited and approved the manuscript. R.M. serves as the author responsible for contact and ensures communication.

## Abstract

Sensing of heat, high light (HL), or mechanical injury by a single leaf of a plant results in the activation of different systemic signals that reach systemic tissues within minutes and trigger systemic acquired acclimation (SAA) or systemic wound responses (SWRs), resulting in a heightened state of stress readiness of the entire plant. Among the different signals associated with rapid systemic responses to stress in plants are electric, calcium and reactive oxygen species (ROS) waves. These signals propagate from the stressed or injured leaf to the rest of the plant through the plant vascular bundles, and trigger SWRs and SAA in systemic tissues. However, whether they can propagate through other cell types, and whether or not they are interlinked, remain open questions. Here we report that in response to wounding or heat stress (HS), but not HL stress, the ROS wave can propagate through mesophyll cells of *Arabidopsis thaliana*. Moreover, we show that propagation of the ROS wave through mesophyll cells during these stresses is sufficient to restore SWR and SAA transcript accumulation in systemic leaves, as well as SAA to HS (but not HL). We further show that propagation of the ROS wave through mesophyll cells could contribute to systemic signal integration during HL&HS stress combination. Our findings reveal that the ROS wave can propagate through tissues other than the vascular bundles of plants, and that different stresses can trigger different types of systemic signals that propagate through different cell layers and induce stress-specific systemic responses.

**One-sentence summary:** In addition to vascular bundles, mesophyll cells can mediate the ROS wave during systemic responses to wounding or heat stress in Arabidopsis.

## INTRODUCTION

In response to different abiotic stresses plants mount an acclimation response that counters the adverse effects of stress on plant metabolism, reproduction, and overall survival (Zhu, 2016; Kollist et al., 2019; Cheung et al., 2020). This response is triggered upon perception of stress at the tissues immediately subjected to stress (termed local tissues), as well as in other tissues of the plant that have not yet experienced the stress (termed systemic tissues). The perception of stress at the local tissues activates therefore a signal transduction process that links the different tissues (local to all systemic tissues) over long distances, sometime spanning the entire length of the plant (*e.g.,* Miller et al., 2009; Szechyńska-Hebda et al., 2010; Christmann et al., 2013; Choi et al., 2014; Gilroy et al., 2016; Guo et al., 2016; Choi et al., 2017; Choudhury et al., 2018; Devireddy et al., 2018; Takahashi et al., 2018; Toyota et al., 2018; Fichman et al., 2019; Wang et al., 2019; Zandalinas et al., 2019; Devireddy et al., 2020b; Devireddy et al., 2020a; Farmer et al., 2020; Fichman and Mittler, 2020). This process is termed systemic signaling, and the acclimation of systemic tissues to stress, upon perception of the systemic signal, is called systemic acquired acclimation (SAA; Karpinski et al., 1999). A similar systemic signaling process occurs in plants upon wounding, and this process is termed systemic wound response (SWR; Walker-Simmons et al., 1984). During SAA or SWR, different abiotic stress- or wound-response transcripts and hormones that rapidly accumulate in the local leaf upon stress or wounding also accumulate within minutes in the systemic tissues, and these transcripts and hormones are thought to mediate SAA or SWR at the systemic tissues (*e.g.,* Galvez-Valdivieso et al., 2009; Miller et al., 2009; Suzuki et al., 2013; Zandalinas et al., 2019; Fichman et al., 2020b). Although the process of SAA or SWR can be easily traced back to some of the regulatory transcripts and hormones that accumulate in systemic tissues during stress, how the systemic signal initiating at the local leaf and reaching the systemic tissues is propagated, and what is its nature, are still ongoing subjects of active research (*e.g.,* Fichman et al., 2020a; Fichman and Mittler, 2020). Among the main candidates for systemic signals mediating SAA or SWRs are electric, calcium, reactive oxygen species (ROS), and hydraulic pressure waves (Miller et al., 2009; Christmann et al., 2013; Mousavi et al., 2013; Choi et al., 2014; Nguyen et al., 2018; Toyota et al., 2018; Shao et al., 2020). Recently, wound-induced electric and calcium waves were shown to be dependent on the function of glutamate receptor-like (GLR) calcium channels expressed at the vascular bundles of *Arabidopsis thaliana*, and a double mutant for *glr3.3;glr3.6* was shown to be deficient in wound-induced systemic signaling (Mousavi et al., 2013; Nguyen et al., 2018; Toyota et al., 2018; Shao et al., 2020). In contrast, systemic responses, and SAA to high light (HL) or heat stress (HS) were found to be dependent on the ROS wave and mediated by the respiratory burst oxidase homolog proteins D/F (RBOHD and/or RBOHF; Miller et al., 2009; Fichman et al., 2019; Zandalinas et al., 2020b). At least in response to HL stress, this process was also found to occur at the vascular bundles of Arabidopsis (Zandalinas et al., 2020b). A new study has now revealed that GLR3.3 and/or GLR3.6 are not absolutely required for HL-induced systemic ROS signaling, and that the systemic signal mediating SAA to HL stress in Arabidopsis requires a coordinated function of plasmodesmata (PD) proteins (*i.e.,* plasmodesmata-localized proteins 1 and 5; PDLP1 and PDLP5) and RBOHD (Fichman et al., 2021). In addition, aquaporins such as PIP2;1 and calcium-permeable channels, such as cyclic nucleotide-gated calcium channel 2 (CNGC2), and mechanosensitive small conductance–like (MSL) channels 2 and 10 were found to be involved in this process (Fichman et al., 2021). Moreover, in response to wounding the ROS wave was shown to induce a redox wave that propagated throughout the plant within minutes (Fichman and Mittler, 2021).

A recent study has also revealed that in contrast to the local application of HL or HS to a single leaf of Arabidopsis, or the co-application of HL and HS to the same leaf (HL+HS), the co-application of HL and HS to two different leaves of the same plant (HL&HS) resulted in a stronger ROS wave response (Zandalinas et al., 2020a). It was further found that the plant hormone jasmonic acid (JA) suppresses the activation of the ROS wave in local leaves simultaneously subjected to a combination of HL and HS (HL+HS; Zandalinas et al., 2020a). Although the ROS wave was found to propagate through the vascular bundles of Arabidopsis during systemic responses to HL stress (Zandalinas et al., 2020b), it is unknown at present whether it propagates through the same plant tissues during other stresses, such as HS or wounding. Finding, for example, that the ROS wave propagates through other plant tissues during HS, could provide a potential explanation to the stronger ROS wave signal observed under conditions of HL&HS (Zandalinas et al., 2020a). In addition, it could provide initial evidence for the propagation of rapid systemic signals outside the vascular bundles of plants.

To identify the plant tissues that mediate RBOHD-dependent systemic ROS signal propagation during responses to HS or wounding, we used the *rbohD* transgenic lines we previously developed to study the propagation of the ROS wave during HL stress (Zandalinas et al., 2020b). Our findings reveal that in contrast to RBOHD-dependent systemic responses to HL stress, that were exclusively mediated through the vascular bundles of Arabidopsis (Zandalinas et al., 2020b), RBOHD-dependent systemic signaling during HS (Zandalinas et al., 2020a), or wounding (Miller et al., 2009; Fichman et al., 2019; Fichman and Mittler, 2021), are mediated through the vascular bundles and/or mesophyll cells. We further show that propagation of the ROS wave through mesophyll cells could contribute to the stronger systemic ROS signal observed in plants subjected to HL and HS simultaneously applied to two different leaves (HL&HS; Zandalinas et al., 2020a). Our findings provide direct evidence for the propagation of rapid systemic ROS signals through tissues other than the vascular bundles of Arabidopsis.

## RESULTS

### Vascular bundles or mesophyll cells can mediate the ROS wave during the systemic response of Arabidopsis to wounding

To identify the plant tissues that transmit RBOHD-dependent systemic signaling (*i.e.,* the ROS wave; Miller et al., 2009) in response to a local application of wounding, we used the different transgenic lines we previously developed of *rbohD*, in which *RBOHD* was expressed under its native promoter or different tissue-specific promoters (Zandalinas et al., 2020b). These lines were previously characterized for their ROS wave propagation, SAA and *Zat12* expression in response to a local application of HL stress, and the localization and stable expression of the RBOHD protein in their different tissues was confirmed using GFP-RBOHD fusions (Zandalinas et al., 2020b). In our analysis we included wild-type, *rbohD* null mutants, and *rbohD* mutants in which *RBOHD* was expressed under its native promoter, or epidermis-, mesophyll- xylem parenchyma-, phloem- or bundles sheath-specific promoters (Zandalinas et al., 2020b). All plants were wounded on a single local leaf and local and systemic ROS levels were imaged in whole-plants grown in soil using the new live-imaging method we developed to image ROS (Fichman et al., 2019; Zandalinas et al., 2020a; Zandalinas et al., 2020b). As shown in Figure 1, wound-induced systemic ROS accumulation was suppressed in the *rbohD* mutant and this suppression was complemented to wild type levels by expression of *RBOHD* in the *rbohD* mutant using its native promoter. Expressing the RBOHD protein in *rbohD* plants using the mesophyll-, xylem parenchyma- or phloem-specific promoters also complemented the systemic accumulation of ROS to wild type levels in the *rbohD* mutant in response to wounding. In contrast, as shown in Supplementary Figure S1, as well as previously reported (Zandalinas et al., 2020b), in response to a local application of HL stress, expression of the RBOHD protein in *rbohD* plants using the native promoter of *RBOHD*, or using the xylem parenchyma- or phloem-specific promoters (but not the mesophyll-specific promoter), complemented the systemic accumulation of ROS in *rbohD* mutants to wild type levels in response to HL. These finding reveal that in response to a local wounding treatment, the ROS wave can propagate through the vascular bundles, or mesophyll cells of Arabidopsis.

**Figure 1.**
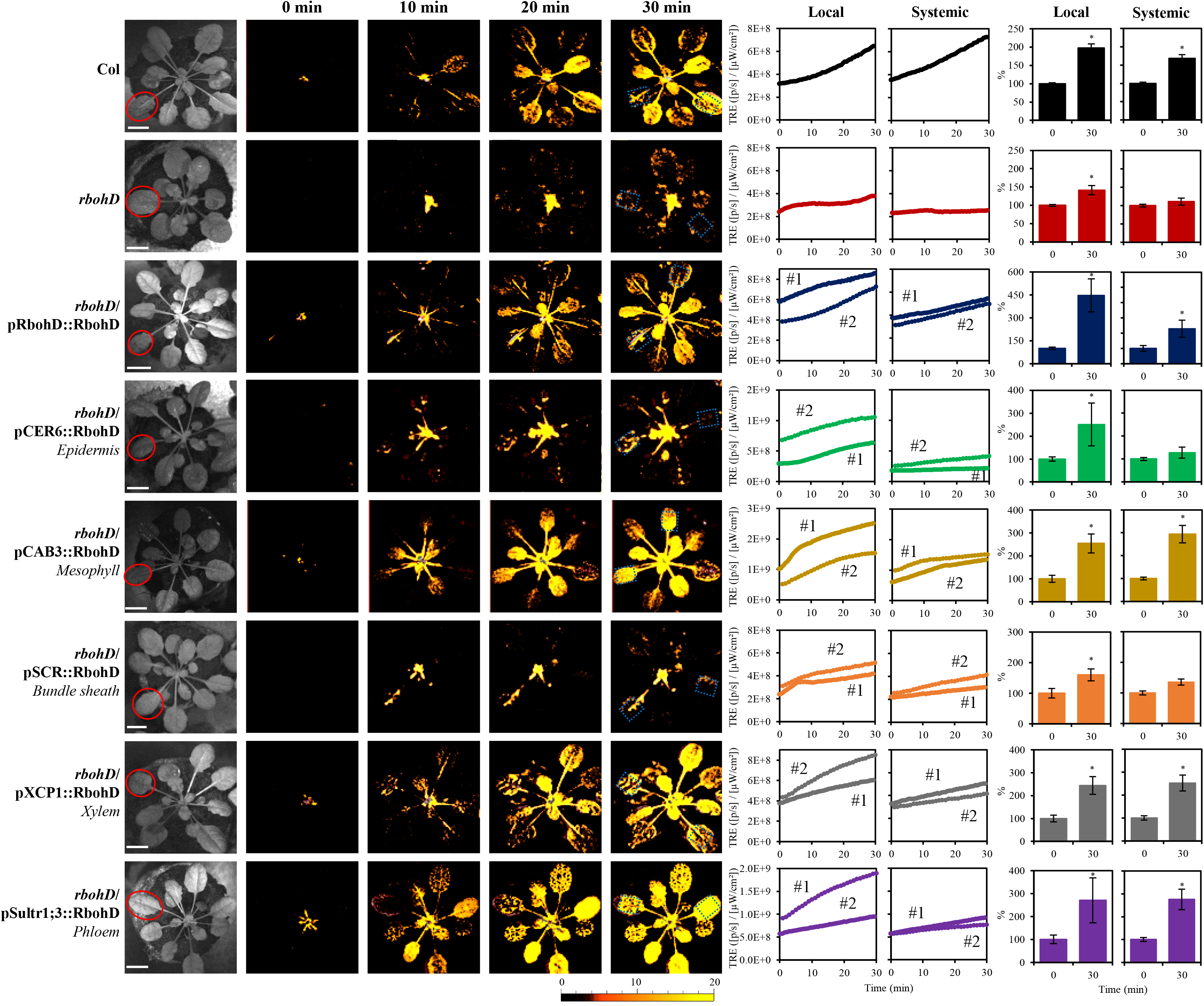
Complementation of wound-induced local and systemic ROS signaling in the *rbohD* mutant with *RBOHD* driven by different tissue-specific promoters. Representative time-lapse images of whole-plant ROS levels in wild type, *rbohD* and the different *rbohD*-complemented *Arabidopsis thaliana* plants subjected to a local wound treatment (red circles), are shown on left; representative line graphs showing continuous measurements of ROS levels in local and systemic leaves of wild type, *rbohD*, and two independent homozygous complemented lines (#1 and #2), over the entire course of the experiment (0 to 30 min) are shown in the middle (ROIs for some of them are indicated with blue boxes); and statistical analysis of ROS levels in local and systemic leaves at 0 and 30 min is shown on right (Student t-test, SD, N=10, *p < 0.05). All experiments were repeated at least 3 times with similar results. Scale bar indicates 1 cm. *Abbreviations used*: RBOHD, respiratory burst oxidase homolog D; CER, eceriferum; CAB, chlorophyll A/B binding protein; SCR, scarecrow; XCP, xylem cysteine peptidase; ROI, region of interest; Sultr, sulfate transporter; TRE, total radiant efficiency.

### Vascular bundles or mesophyll cells can mediate the ROS wave during the systemic response of Arabidopsis to heat stress

As shown in Figure 2, a similar result to that shown in Figure 1 was obtained when a local Arabidopsis leaf was subjected to HS. Thus, similar to the local application of wounding (Figure 1), but different from the local application of HL (Supplementary Figure S1; Zandalinas et al., 2020b), expression of the RBOHD protein in *rbohD* plants using its native promoter, or using the mesophyll-, xylem parenchyma- or phloem-specific promoters, complemented the systemic accumulation of ROS in *rbohD* mutants to wild type levels in response to a local application of HS. The findings shown in Figures 1 and 2 reveal therefore that unlike rapid systemic ROS responses to HL, that could only be complemented to wild type levels in the *rbohD* mutant by expressing the RBOHD protein in xylem parenchyma or phloem cells (Supplementary Figure S1; Zandalinas et al., 2020b), tissues limited in their localization to the vascular bundles, systemic ROS responses (*i.e.,* the ROS wave) to wounding or HS can be mediated by RBOHD protein found in mesophyll cells that are primarily localized outside the vascular bundles of plants.

**Figure 2.**
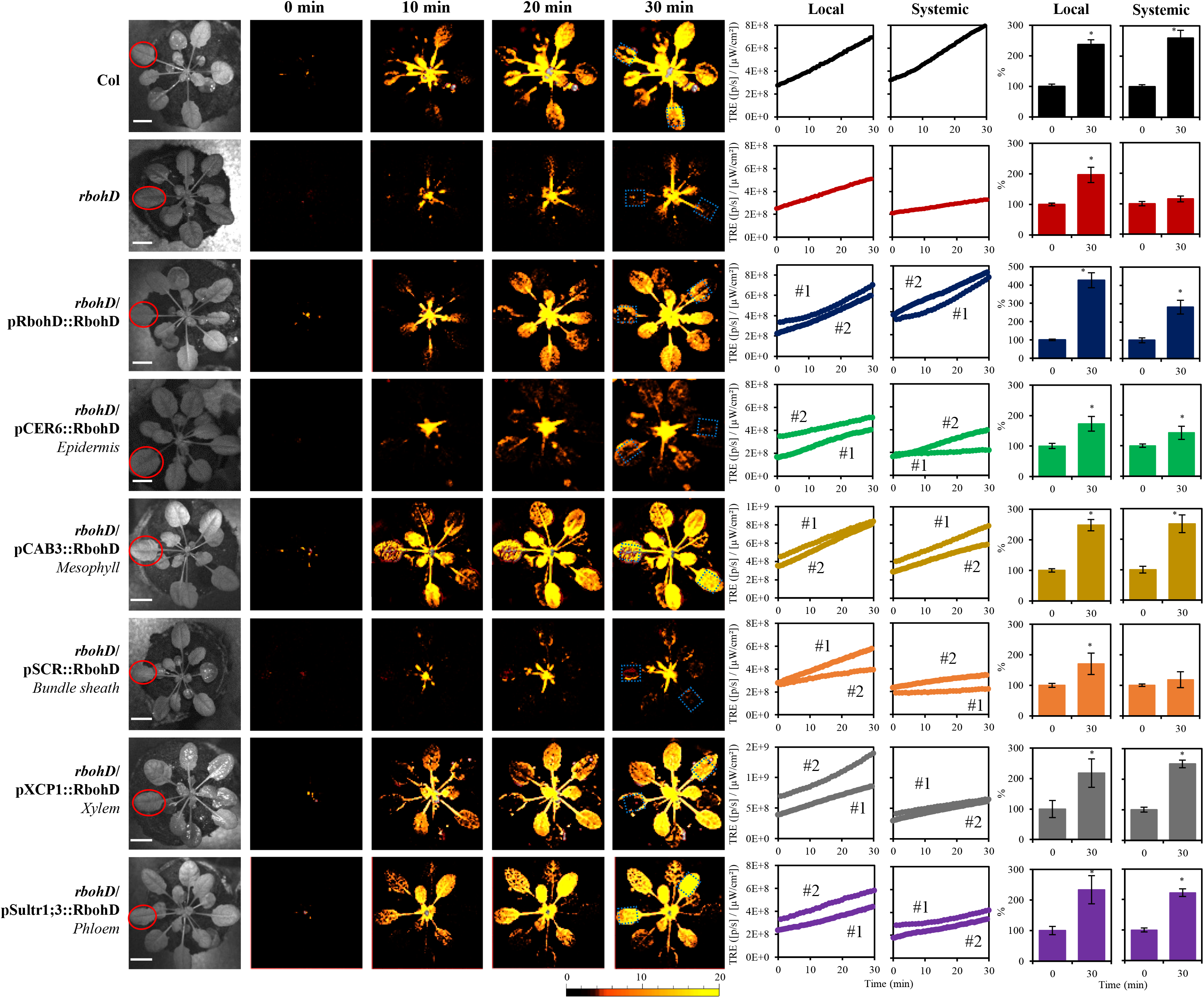
Complementation of heat stress-induced local and systemic ROS signaling in the *rbohD* mutant with *RBOHD* driven by different tissue-specific promoters. Representative time-lapse images of whole-plant ROS levels in wild type, *rbohD* and the different *rbohD*-complemented *Arabidopsis thaliana* plants subjected to a local heat stress treatment (red circles), are shown on left; representative line graphs showing continuous measurements of ROS levels in local and systemic leaves of wild type, *rbohD* and two independent homozygous complemented lines (#1 and #2), over the entire course of the experiment (0 to 30 min) are shown in the middle (ROIs for some of them are indicated with blue boxes); and statistical analysis of ROS levels in local and systemic leaves at 0 and 30 min is shown on right (Student t-test, SD, N=10, *p < 0.05). All experiments were repeated at least 3 times with similar results. Scale bar indicates 1 cm. *Abbreviations used*: RBOHD, respiratory burst oxidase homolog D; CER, eceriferum; CAB, chlorophyll A/B binding protein; SCR, scarecrow; XCP, xylem cysteine peptidase; ROI, region of interest; Sultr, sulfate transporter; TRE, total radiant efficiency.

### Complementing the ROS wave by expression of RBOHD in mesophyll cells restores SAA- and SWR-associated transcript expression in systemic leaves in response to a local HS or wounding treatment

Complementing the ROS wave by expression of *RBOHD* in mesophyll cells (Figures 1 and 2) might or might not complement the expression of systemic transcripts previously associated with SAA or SWR in response to a local application of HS or wounding, respectively. Complementation of *RBOHD* expression in the *rbohD* mutant using the xylem parenchyma- or phloem- (but not mesophyll-) specific promoters restored the expression of the *Zat12* SAA and SWR gene in response to local application of HL stress (measured using Zat12:luciferase;*rbohD* double mutants complemented with the different tissue-specific *RBOHD* transformation vectors; Zandalinas et al., 2020b). Because Zat12 reporter plants might not be a good experimental tool to study stress-specific responses to HS, HL or wounding (*Zat12* is expressed in response to HL or wounding; Miller et al., 2009), we elected to study the expression of different wounding-, HS-, or HL-specific transcripts in the different lines shown in Figures 1 and 2 in response to a local application or HS, wounding, or HL using quantitative RT-PCR (qPCR). We chose transcripts for this analysis based on our previous RNA-Seq studies of systemic signaling in response to HL and/or HS (Suzuki et al., 2013; Zandalinas et al., 2019; Fichman et al., 2020b; Zandalinas et al., 2020a), as well as based on studies of systemic wound responses using transcriptomics and qPCR analyses (Suzuki et al., 2013; Toyota et al., 2018). As shown in Figure 3A, expression of the wound-response transcripts *JAZ5* and *JAZ7* was enhanced in local and systemic leaves of wild type plants upon local wounding. In contrast, in response to the same treatment, the expression of these transcripts was suppressed in systemic (but not local) leaves or the *rbohD* mutant. Complementation of *RBOHD* expression with the *RBOHD* native promoter, or the mesophyll-, xylem parenchyma-, or phloem-specific promoters restored the systemic expression of *JAZ5* and *JAZ7* in response to a local wounding treatment. In contrast, complementation of *RBOHD* expression with the bundle sheath- or epidermis-specific promoters failed to restore the systemic expression of *JAZ5* and *JAZ7* to wild type levels in response to the local wounding treatment. These findings reveal that complementing the ROS wave by expression of RBOHD in mesophyll, xylem parenchyma or phloem cells of the *rbohD* mutant was sufficient to restore some SWR-specific transcript expression in response to a local wounding treatment.

**Figure 3.**
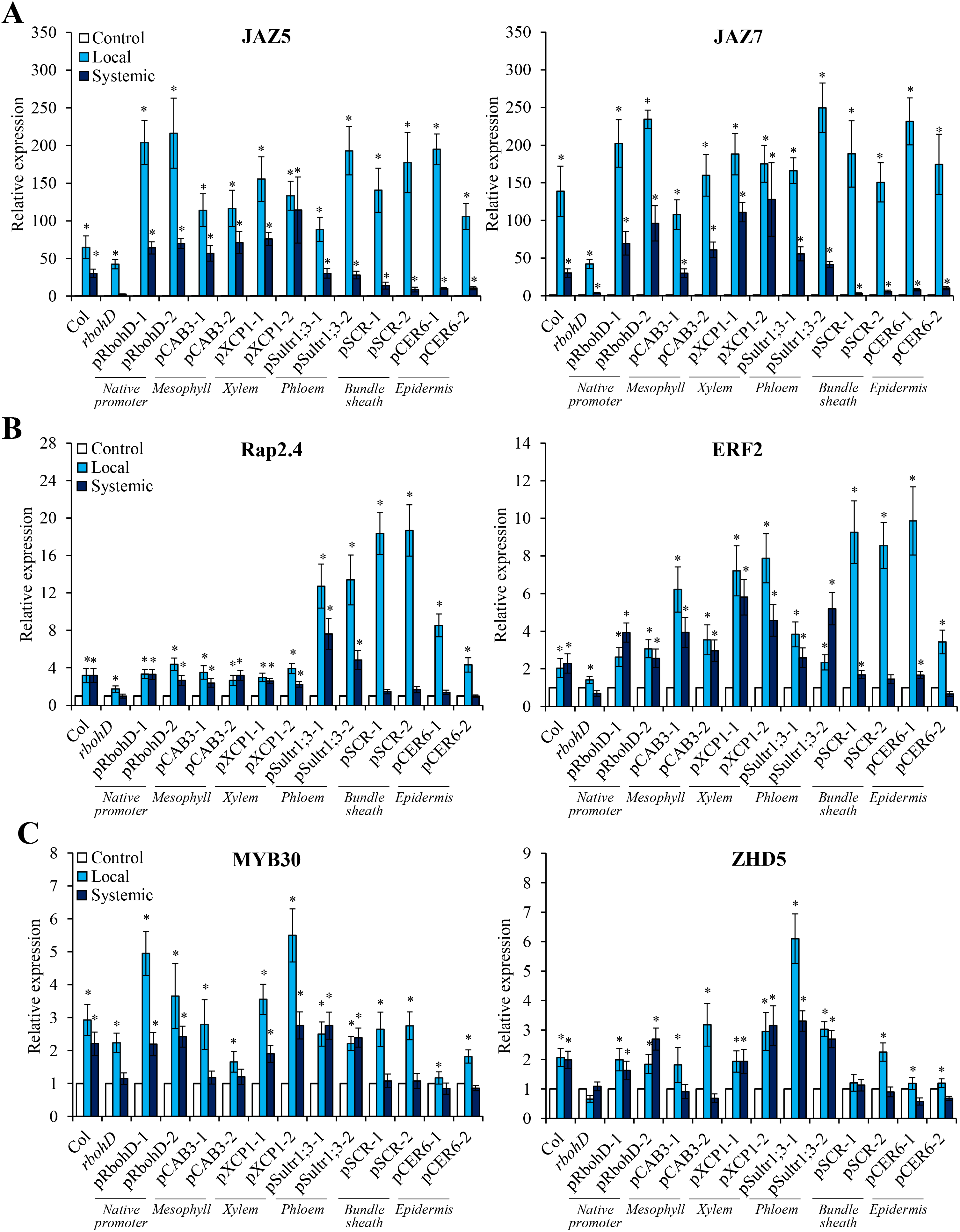
Local- and systemic stress-induced transcript expression in wild type, *rbohD*, and the *rbohD* mutant complemented with *RBOHD* driven by different tissue-specific promoters. (A) Local and systemic steady-state levels of *JAZ5* (AT1G17380) and *JAZ7* (AT2G34600) transcripts in wild type, *rbohD*, and the different *rbohD*-complemented *Arabidopsis thaliana* plants subjected to a local wound treatment. (B) Local and systemic steady-state levels of *Rap2.4* (AT1G78080) and *ERF2* (AT5G47220) transcripts in wild type, *rbohD*, and the different *rbohD*-complemented Arabidopsis plants subjected to a local heat-stress treatment. (C) Local and systemic steady-state levels of *MYB30* (AT3G28910) and *ZHD5* (AT1G75240) transcripts in wild type, *rbohD*, and the different *rbohD*-complemented Arabidopsis plants subjected to a local high light-stress treatment. Student t-test, SD, N=3, *p < 0.05. *Abbreviations used*: RBOHD, respiratory burst oxidase homolog D; CER, eceriferum; CAB, chlorophyll A/B binding protein; SCR, scarecrow; XCP, xylem cysteine peptidase; Sultr, sulfate transporter; JAZ, jasmonate-zim-domain protein; ERF, ethylene response factor; ZHD, zinc-finger homeodomain.

To test the effect of restoring *RBOHD* expression in the different tissues on SAA responses to HS, we studied the expression of *Rap2.4* and *ERF2*, two transcripts previously associated with SAA to HS (Suzuki et al., 2013; Zandalinas et al., 2020a), in local and systemic leaves of the different wild type, *rbohD* and *rbohD*-complemented lines, in response to a local HS treatment. As shown in Figure 3B, expression of the HS-response transcripts *Rap2.4* and *ERF2* was enhanced in local and systemic leaves of wild type plants upon a local HS treatment. In contrast, in response to the same treatment, the enhanced expression of these transcripts was blocked in systemic and suppressed in local leaves or the *rbohD* mutant. Complementation of *RBOHD* expression with the *RBOHD* native promoter, or the mesophyll-, xylem parenchyma-, or phloem-specific promoters restored the systemic expression of *Rap2.4* and *ERF2* in response to a local HS treatment. In contrast, complementation of *RBOHD* expression with the bundle sheath- or epidermis-specific promoters did not restore the systemic expression of *Rap2.4 and ERF2* to wild type levels in response to a local HS treatment. These findings reveal that similar to the response of *JAZ5* and *JAZ7* to wounding (Figure 3A), restoring the ROS wave by expression of *RBOHD* in mesophyll, xylem parenchyma or phloem cells was sufficient to restore some SAA transcript expression in systemic leaves in response to a local HS treatment.

To study whether a similar effect would occur in complemented *rbohD* plants subjected to a local treatment of HL, we studied the expression of *MYB30* and *ZHD5*, two transcripts associated with the SAA response of Arabidopsis to HL stress (Zandalinas et al., 2019; Fichman et al., 2020b; Zandalinas et al., 2020a). As shown in Figure 3C, similar to the complementation of *Zat12* expression in the different *rbohD*-complemented lines (Zandalinas et al., 2020b), complementation of *RBOHD* expression in xylem parenchyma or phloem (but not mesophyll) cells of the *rbohD* mutant supported the systemic expression of *MYB30* and *ZHD5* in response to a local treatment of HL stress. Taken together, the results presented in Figures 1–3 and Supplementary Figure S1 reveal that complementing the expression of *RBOHD* in the mesophyll, xylem parenchyma or phloem cells of the *rbohD* mutant restores not only the ROS wave, but also the expression of certain systemic transcripts specific to wounding or HS. In contrast, complementing the expression of *RBOHD* in mesophyll cells of the *rbohD* mutant did not complement the ROS wave or systemic HL-specific SAA transcripts in response to a local application of HL stress.

### Complementing the ROS wave by expression of *RBOHD* in mesophyll cells restores local HS-induced SAA

Complementing the expression of *RBOHD* in the xylem parenchyma or phloem cells of the *rbohD* mutant restored SAA to HL (Supplementary Figure S2; Zandalinas et al., 2020b). Although we do not have a biological assay for SAA during SWR, aside from measuring systemic wound-induced transcript expression as shown in Figure 3, an assay for SAA to HS was previously reported (Suzuki et al., 2013; Zandalinas et al., 2020a). We therefore used this assay to study whether restoring *RBOHD* expression in mesophyll cells could restore SAA to HS of the *rbohD* mutant. As shown in Figure 4, complementing the expression of *RBOHD* in the *rbohD* mutant using its native promoter, or the mesophyll-, xylem parenchyma-, or phloem-specific promoters restored SAA to HS. In contrast, complementing the expression of *RBOHD* in the *rbohD* mutant using the mesophyll-specific promoter failed to restore SAA to HL (Supplementary Figure S2; Zandalinas et al., 2020b). The findings presented in Figures 1–4 and Supplementary Figures S1 and S2 reveal therefore that expression of *RBOHD* in mesophyll cells can restore the ROS wave, systemic transcript expression and SAA to HS (but not HL stress) in the *rbohD* mutant.

**Figure 4.**
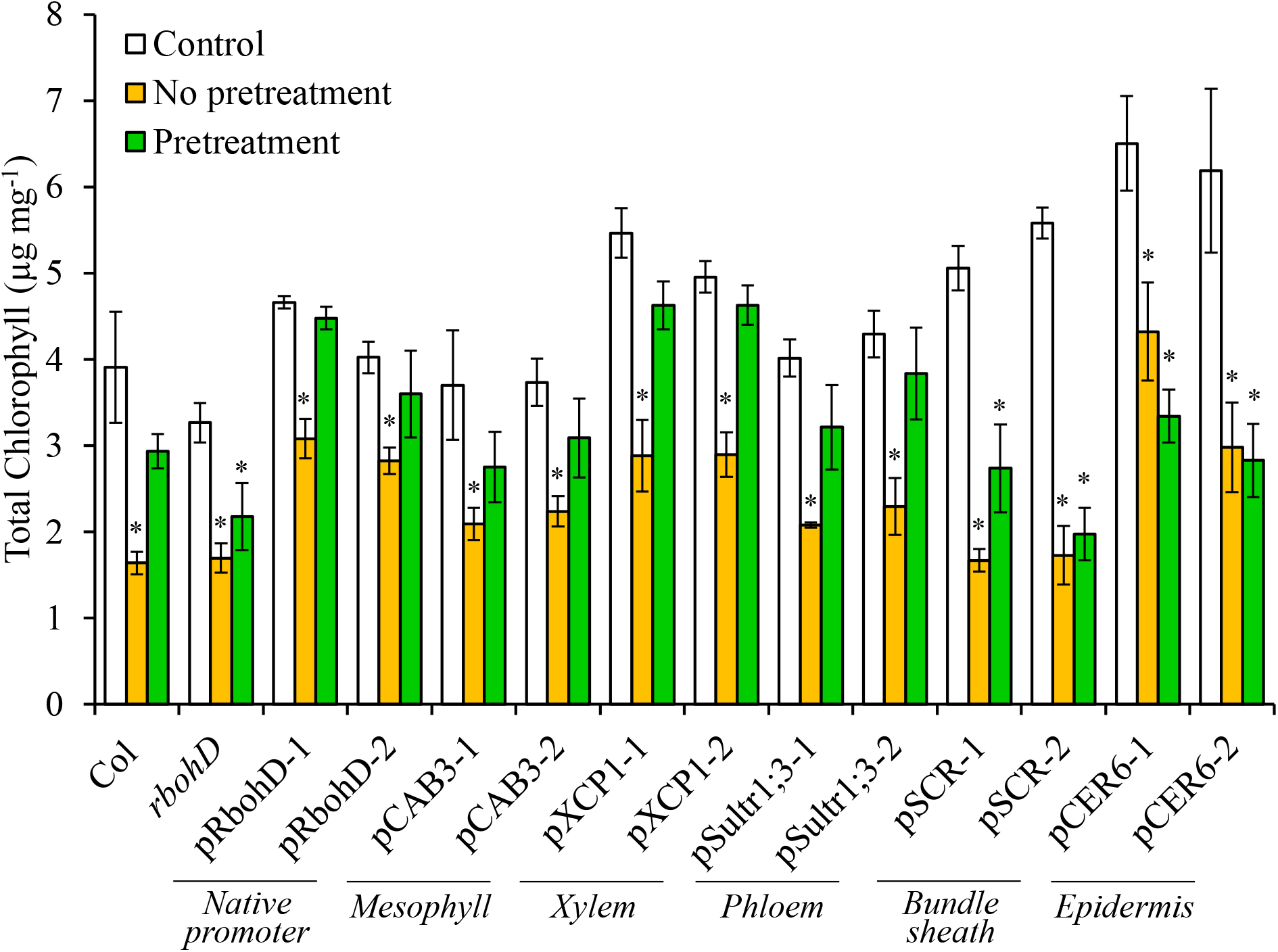
Complementation of heat stress-induced SAA in the *rbohD* mutant with *RBOHD* driven by different tissue-specific promoters. Heat stress-induced changes in systemic leaf chlorophyll content of wild type, *rbohD* and the different *rbohD*-complemented *Arabidopsis thaliana* plants are shown. Systemic leaves were obtained from plants that were either untreated and unstressed (Control), untreated at their local leaves and subjected to a systemic heat-stress treatment (No pretreatment), or subjected to a local heat stress pre-treatment before being subjected to a systemic heat stress treatment (Pretreatment). SAA is evident when the systemic leaf of a pre-treated plant does not show a loss of chlorophyll content following a systemic heat stress treatment. Ten different plants each from two independent complemented lines for each construct were subjected to the SAA heat stress assay and chlorophyll content was measured in systemic leaves. Student t-test, SD, N=10, *p < 0.05. *Abbreviations used*: RBOHD, respiratory burst oxidase homolog D; CER, eceriferum; CAB, chlorophyll A/B binding protein; SCR, scarecrow; XCP, xylem cysteine peptidase; Sultr, sulfate transporter.

### Could expression of *RBOHD* in mesophyll cells contribute to the stronger systemic ROS signal observed in plants subjected to HL&HS?

We previously reported that HS and HL, when applied to two different leaves of the same Arabidopsis plant (HL&HS), result in a stronger ROS wave response compared to HS or HL applied to a single leaf, or to the same leaf (HL+HS; Zandalinas et al., 2020a). Our current findings that in response to HS the ROS wave could be mediated through mesophyll, xylem parenchyma, and/or phloem cells (Figures 2–4), but in response to HL it could only be mediated through xylem parenchyma and/or phloem cells (Supplementary Figures S1 and S2; Zandalinas et al., 2020b), might provide a potential explanation to this phenomena. In response to HL and HS applied to two different leaves (HL&HS), the systemic ROS wave might be stronger because it would propagate through an additional cell layer (mesophyll, contributed by the HS treatment). This could not occur of course when the two stresses are applied to the same leaf because under these conditions the ROS wave induced by HL+HS applied to the same leaf is suppressed by JA (Zandalinas et al., 2020a). To test whether the ROS wave could propagate through mesophyll cell layers during HL&HS combination, we compared the intensity of the ROS wave between wild type, *rbohD*, and *rbohD* in which *RBOHD* expression was complemented at the mesophyll or phloem cells, subjected to a HL&HS treatment (Figure 5 and Supplementary Figure S3). As shown in Figure 5, compared to wild type plants, the ROS wave was suppressed in *rbohD* plants subjected to the HL&HS treatment. Complementation of *RBOHD* expression with *RBOHD* expressed under the control of its native promoter, or a phloem specific promoter (that could restore HS- or HL-response ROS wave functions from the two different leaves; Figures 2–4 and Supplementary Figures S1 and S3; Zandalinas et al., 2020b) restored the ROS wave to its high level of expression. In contrast, complementation of the *rbohD* mutant with *RBOHD* expressed under the mesophyll- specific promoter (that could only restore HS-, but not HL-response ROS wave functions from the HL-treated leaf; Figures 2–4 and Supplementary Figures S1 and S3; Zandalinas et al., 2020b), could not restore the ROS wave to its maximal intensity. These finding demonstrate that under conditions of HL&HS at least part of the ROS wave that spreads throughout the plant (originating from the HS-treated leaf) could be mediated through mesophyll cells. Complementation of *RBOHD* expression with *RBOHD* expressed under the control of the phloem-specific promoter was nonetheless sufficient to restore the ROS wave to wild type or *rbohD* mutant complemented with *RBOHD* under its native promoter levels (Figure 5), suggesting that in wild type plants transmission of the ROS wave signal through phloem cells is sufficient to cause a higher signal during HL&HS combination.

**Figure 5.**
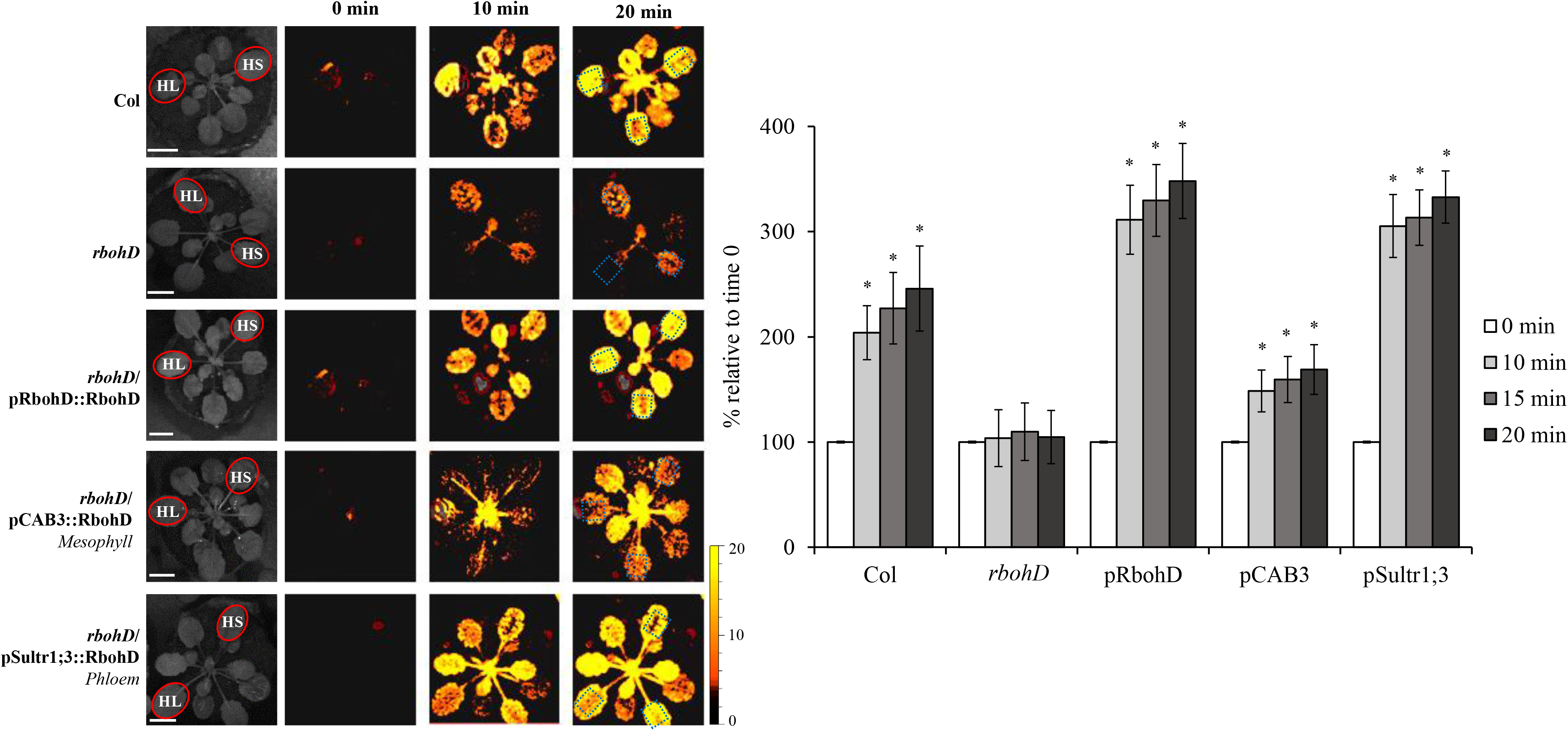
Complementation of light- and heat-induced local and systemic ROS signaling in the *rbohD* mutant with *RBOHD* driven by the phloem- or mesophyll tissue-specific promoters, during stress combination. Representative time-lapse images of whole-plant ROS levels of wild type, *rbohD* and *rbohD*-complemented *Arabidopsis thaliana* plants subjected to a local light (HL) and heat stress (HS) treatments, simultaneously applied to two leaves of the same plant (red circles; HL&HS; Zandalinas et al., 2020a) are shown on left (ROIs for some of them are indicated with blue boxes), and statistical analysis of ROS levels in systemic leaves of treated plants at 0, 10, 15 and 20 min is shown on right (Student t-test, SD, N=10, *p < 0.05). All experiments were repeated at least 3 times with similar results. Scale bar indicates 1 cm. *Abbreviations used*: HL, high light; HS, heat stress; RBOHD, respiratory burst oxidase homolog D; CAB, chlorophyll A/B binding protein; Sultr, sulfate transporter.

## DISCUSSION

Abiotic, mechanical injury, and biotic stresses trigger a rapid systemic signal transduction process that activates different acclimation and defense mechanisms in systemic tissues within minutes of stress sensing at the local tissues (Fichman et al., 2019; Kollist et al., 2019; Fichman and Mittler, 2020). Up until now, the electric, calcium and ROS waves, triggered by wounding or HL stress, were shown to be mediated through the vascular bundles of plants (Mousavi et al., 2013; Nguyen et al., 2018; Toyota et al., 2018; Farmer et al., 2020; Shao et al., 2020; Zandalinas et al., 2020b). Here, we present evidence that in addition to vascular bundles, mesophyll cells can also mediate the ROS wave in response to a local treatment of wounding or HS (Figures 1–6). Mesophyll cells are not typically considered part of the vascular bundles of plants and could occur within leaves and stems as cell layers that connect the vascular tissues to the epidermis, stomata and/or other leaf/stem structures and cell types. Because the ROS wave propagates from cell-to-cell via mechanisms that require apoplastic and symplastic connectivity between cells (Miller et al., 2009; Fichman et al., 2021), and mesophyll cells are connected with each other via PD and/or their shared apoplastic microenvironment, as well as express *RBOHD* under controlled growth conditions (Zandalinas et al., 2020b), the basic mechanisms that allow the ROS wave to propagate from cell-to-cell through mesophyll cell layers appear to be present. In contrast, GLR3.3 and/or GLR3.6 that are required for rapid wound-response systemic signaling are not thought to be localized to mesophyll cells (they are thought to be exclusively localized to the xylem parenchyma and phloem cells; Mousavi et al., 2013; Nguyen et al., 2018; Toyota et al., 2018; Shao et al., 2020). A recent study has shown that GLR3.3 and/or GLR3.6 are not absolutely required for the ROS wave to propagate in response to a local treatment of HL stress (Fichman et al., 2021). Taking this study into consideration, it is plausible that the ROS wave will propagate through tissues that do not express GLR 3.3 and/or GLR3.6, possibly using other calcium-permeable channels such as CNGCs or MSLs (Fichman et al., 2021). Having many of the required proteins and physical connections/proximity required for ROS cell-to-cell signals to occur, support the possibility that the ROS wave can propagate through layers of mesophyll cells that are outside the vascular bundles (Figure 6).

**Figure 6.**
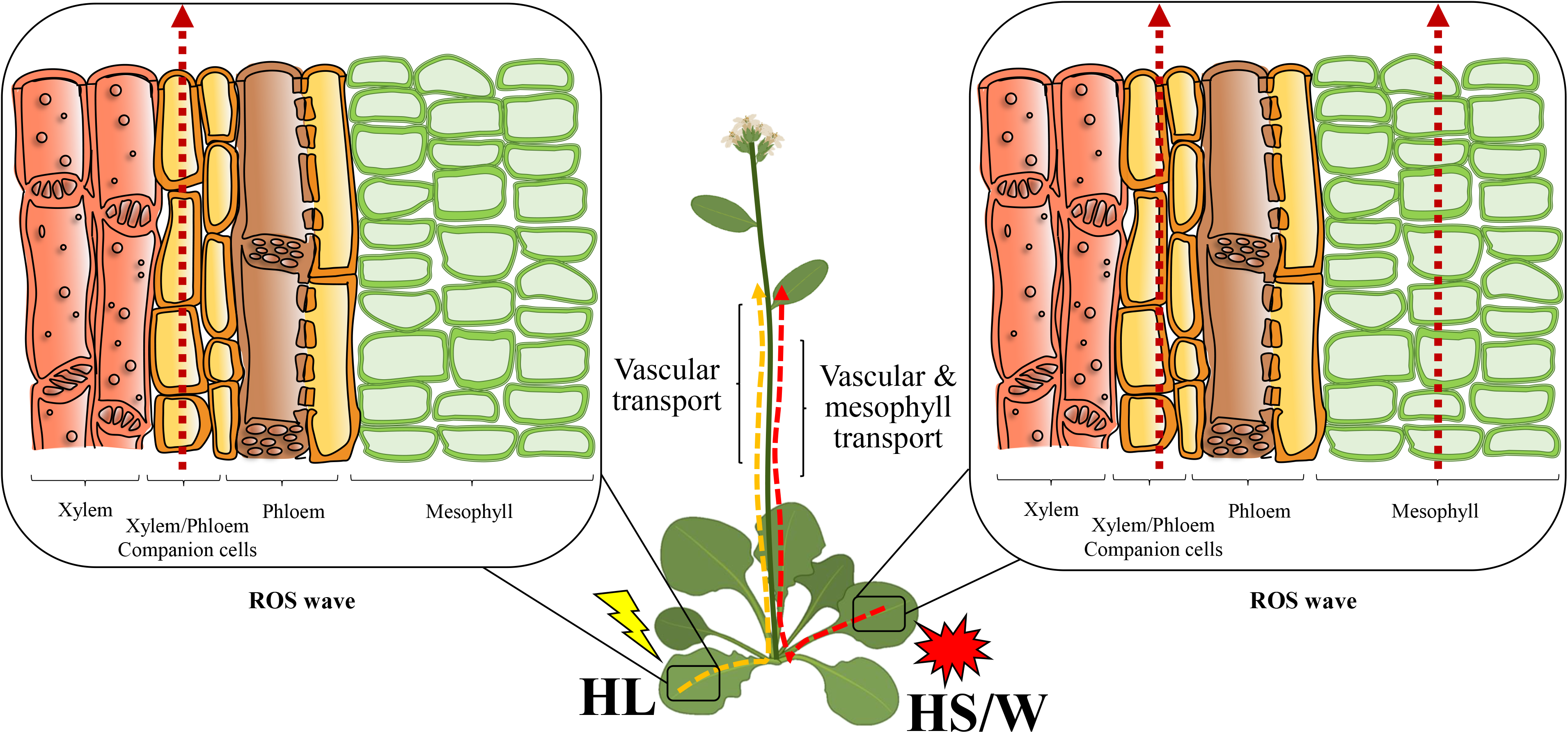
A model showing that when light stress is applied to a local leaf, the ROS wave is mediated through vascular bundles. In contrast, when heat stress or wounding are applied to a local leaf, both vascular and mesophyll cells can mediate the ROS wave. *Abbreviations used*: HL, high light; HS, heat stress; W, wounding; ROS, reactive oxygen species.

In addition to showing that the ROS wave can propagate outside the vascular bundles of Arabidopsis (Figures 1 and 2), supporting systemic wound- and HS-induced transcript expression in systemic leaves (Figure 3) and mediating SAA to HS (Figure 4), our findings further highlight the interesting possibility that different stresses, *e.g.,* HS, HL and wounding, trigger different types of systemic waves that propagate through different tissues, and could even be spatially separated from each other. For example, complementing *RBOHD* expression in mesophyll cells of the *rbohD* mutant can complement systemic responses to wounding (Figure 3). Under these conditions, the electric and calcium waves could propagate through the vascular bundles (supported by GLR3.3;GLR3.6; Mousavi et al., 2013; Nguyen et al., 2018; Toyota et al., 2018; Shao et al., 2020), while the ROS wave could propagate through mesophyll cells (supported by RBOHD; Figures 1 and 3; Zandalinas et al., 2020b). This possibility suggests that the ROS wave can be spatially separated from the calcium and electric waves. Different stresses could therefore trigger different combinations of waves that could travel through different tissues and cell layers of the plant. During systemic responses to HL stress however the separation of systemic signals cannot occur (for reasons unknown at present), and the ROS wave must propagate together with the electric and calcium waves through the vascular bundles. Further studies are of course needed to address these intriguing possibilities.

Under all stresses studied here (HL, HS, wounding), RBOHD appeared to be required for systemic transcript accumulation (Figure 3), suggesting that even though GLRs were present and most likely functional in the *rbohD* mutant, they could not mediate their function to drive the expression of systemic transcript accumulation in the absence of the ROS wave. The ROS wave, even occurring at tissues other than the vascular bundles (*i.e.,* mesophyll cell layers; Figures 1 and 2) could therefore be required to support other systemic signal propagation (such as electric and calcium waves) occurring at the vascular bundles during HS or wound responses. Although it is unknow at present how changes in ROS at the mesophyll cell layers impact electric and calcium signaling at the vascular bundles, one intriguing possibility is that different metabolites, ions, ROS, hormones, and/or pH changes, occurring at the mesophyll cell layers are diffused/transported to the vascular bundles, and these are needed to link the different waves (Fichman et al., 2020a; Fichman and Mittler, 2020). In this respect, it should be mentioned that changes in localized pH levels were recently linked to the triggering and propagation of electric and calcium waves in Arabidopsis (Shao et al., 2020). An alternative explanation could of course be that RBOHF at the vascular tissues replaces the function of RBOHD in linking the different waves during all stresses studied, and that the levels of RBOHF-produced ROS in the vascular bundles of *rbohD* plants complemented by *RBOHD* expressed under the control of a mesophyll-specific promoter are too low to be detected by our assay. Further studies are of course needed to address these possibilities, as well as to resolve the different spatial and temporal relationships that could potentially exist between the different waves, signals and hormones involved in systemic signaling (*e.g.,* Kangasjärvi et al., 2009; Miller et al., 2009; Mittler et al., 2011; Dubiella et al., 2013; Gilroy et al., 2014; Evans et al., 2016; Gilroy et al., 2016; Choi et al., 2017; Fichman et al., 2020a; Fichman and Mittler, 2020).

Our findings that the ROS wave can propagate through multiple cell layers in response to different stresses could also partially explain how the integration of different systemic signals during a combination of HL and HS results in a stronger ROS wave signal (Zandalinas et al., 2020a). It is possible that during a combination of HL and HS applied to two different leaves of the same plant (HL&HS; Zandalinas et al., 2020a), the ROS wave initiated from the two different leaves propagates through all three cell types of the plant (mesophyll, xylem parenchyma and phloem, initiated by the local HS treatment, and xylem parenchyma and phloem, initiated by the local HL treatment). In contrast, during a combination of HL+HS applied to the same leaf, JA suppresses the ROS wave and the signal is lower (Zandalinas et al., 2020a). Our findings that restoring RBOHD expression in mesophyll cells did not result in a stronger systemic ROS signal during a HL&HS treatment (Figure 5), reveals that during HL&HS combination in Arabidopsis the ROS wave could indeed propagate through mesophyll cells (Figures 5 and 6 and Supplementary Figure S3). The ROS wave triggered by the HL treatment (propagating through xylem parenchyma and/or phloem cells) could therefore merge with the ROS wave triggered by the HS treatment (propagating through mesophyll and xylem parenchyma and/or phloem cells) to generate a stronger systemic ROS signal during HL&HS combination that is mediated through multiple cell layers (Figures 5 and 6 and Supplementary Figure S3). Because complementing *RBOHD* expression in the *rbohD* mutant using *RBOHD* expressed under the phloem-specific promoter was sufficient to restore the strong signal observed during HL&HS combination (Zandalinas et al., 2020a; Figure 5, Supplementary Figure S3), it is also possible that the stronger signal observed during HL&HS combination is simply the result of two different ROS wave signals merging together, regardless of the type of tissue supporting their transmission. Further studies are of course needed to dissect the mode of systemic signal integration through the different cell layers during stress combination.

## MATERIALS AND METHODS

### Plant material and growth conditions

*Arabidopsis thaliana* Col-0 (*cv.* Columbia-0), *rbohD* plants (Fichman et al., 2019) and two independent lines each of the different *rbohD* complemented plants (Zandalinas et al., 2020b) were grown in peat pellets (Jiffy-7, Jiffy, http://www.jiffygroup.com/) at 23°C under short day growth conditions (10-hour light/14-hour dark, 50 μmol m^−2^ s^−1^).

### Measurements of ROS accumulation

To image whole-plant ROS levels, plants were fumigated with 50 μM H_2_DCFDA (excitation/emission 495 nm/517 nm; Millipore-Sigma, St. Louis, MO, USA) in 50 mM phosphate buffer (pH 7.4) containing 0.01% Silwet L-77 (LEHLE seeds, Round Rock, TX, USA), using a portable mini nebulizer (Punasi Direct, Hong Kong, China) for 30 min as described previously (Fichman et al., 2019; Zandalinas et al., 2020a; Zandalinas et al., 2020b). Following H_2_DCFDA application, local leaves were exposed to wounding, HL stress, HS, or HL and HS applied to two different leaves located at opposite sides of the plant as described by Zandalinas et al., (2020a). Wounding was achieved by puncturing a single leaf with 18 dressmaker pins (Singer, Murfreesboro, TN, USA) as described in (Fichman et al., 2019). Light (HL) stress was applied by subjecting a single leaf to a light intensity of 1700 μmol m^−2^ s^−1^ for 2 min using a ColdVision fiber optic LED light source (Schott A20980, Southbridge, MA, USA) as described in (Devireddy et al., 2018; Zandalinas et al., 2019; Zandalinas et al., 2020a; Zandalinas et al., 2020b). Heat stress (HS) was induced by placing a heat block 2 cm underneath the treated leaf for 2 min, increasing the leaf temperature to 31-33°C (Zandalinas et al., 2020a). Local and systemic leaf temperature were measured under all conditions and treatment using an infrared camera (C2; FLIR Systems) as previously described (Zandalinas et al., 2020a). Imaging of ROS accumulation in response to a local stress treatment was conducted with an IVIS Lumina S5 platform using Living Image 4.7.2 software (PerkinElmer) as described in (Fichman et al., 2019; Zandalinas et al., 2020a; Zandalinas et al., 2020b). All experiments were repeated at least three times each with 10 wild type, *rbohD* and the different *rbohD* complemented plants.

### RT-qPCR analysis

To analyze transcript expression by RT-qPCR, plants were subjected to a local treatment of wounding, 8-min HL or 8-min HS as described above. Local and systemic leaves were collected and immediately frozen in liquid nitrogen following the 8-min HL or HS treatments, or 30 min following wounding. Relative expression analysis by RT-qPCR was performed according to (Balfagón et al., 2019) by using the CFX Connect Real-Time PCR Detection System (Bio-Rad) and gene-specific primers (Supplementary Table S1; Primer efficiency range of 0.99-1.04). All experiments were repeated at least three times each with at least 5 wild type, *rbohD* and the different *rbohD* complemented plants.

### Heat stress acclimation assay

For heat stress acclimation, a single leaf was pre-treated for 15 min at 31-33°C by placing a heat block 2 cm underneath the treated leaf (Zandalinas et al., 2020a). Plants were then incubated for 45 minutes under controlled conditions. Following the recovery period, a systemic leaf of pre-treated and untreated plants was dipped in a 42°C (or 23°C as control) water bath for 60 min and allowed to recover under controlled growth conditions. Systemic leaves were sampled 6 days after the water bath heat stress treatment for chlorophyll measurements, as previously described (Zandalinas et al., 2020a; Zandalinas et al., 2020c). For HL-induced SAA, a single leaf was pre-treated for 15 min with a light intensity of 1700 μmol m^−2^ sec^−1^ using a ColdVision fiber optic LED light source (Schott A20980, Southbridge, MA, USA). Plants were then incubated for 45 minutes under controlled conditions. Following the recovery period, a systemic leaf was exposed to a light intensity of 1700 μmol m^−2^ sec^−1^ for 45 minutes. Control systemic leaves (untreated) and systemic leaves of plants that were pretreated with HL stress, as described above (SAA), were then analyzed for electrolyte leakage as previously described (Zandalinas et al., 2019; Zandalinas et al., 2020a; Zandalinas et al., 2020b). Acclimation assays were repeated at least 3 times with 10 plants per repeat.

### Statistical analysis

Results are presented as the mean ± SD. Statistical analyses were performed by a two-tailed Student’s t-test (asterisks denote statistical significance at p < 0.05 with respect to controls).

HL: high light
HS: heat stress
RBOHD: respiratory burst oxidase homolog D
ROS: reactive oxygen species
SAA: Systemic acquired acclimation
SWR: systemic wound response

## SUPPLEMENTAL DATA

**Supplementary Table S1.** Transcript-specific primers used for relative expression analysis by RT-qPCR.

**Supplementary Figure S1.** Complementation of light (HL) stress-induced local and systemic ROS signaling in the *rbohD* mutant with *RBOHD* driven by different tissue-specific promoters. Representative time-lapse images of whole-plant ROS levels in wild type, *rbohD* and the different *rbohD* complemented *Arabidopsis thaliana* plants subjected to a local HL-stress treatment (red circles), are shown on left; representative line graphs showing continuous measurements of ROS levels in local and systemic leaves of wild type, *rbohD* and two independent homozygous complemented lines (#1 and #2), over the entire course of the experiment (0 to 30 min) are shown in the middle (ROIs for some of them are indicated with blue boxes); and statistical analysis of ROS levels in local and systemic leaves at 0 and 30 min is shown on right (Student t-test, SD, N=10, *p < 0.05). All experiments were repeated at least 3 times with similar results. Scale bar indicates 1 cm. *Abbreviations used*: HL, high light; RBOHD, respiratory burst oxidase homolog D; CER, eceriferum; CAB, chlorophyll A/B binding protein; SCR, scarecrow; XCP, xylem cysteine peptidase; ROI, region of interest; Sultr, sulfate transporter; TRE, total radiant efficiency. The experiments shown were conducted in parallel to the experiments shown in Figures 1 and 2 and are a repeat of the study reported previously (Zandalinas et al., 2020b), with similar results.

**Supplementary Figure S2.** Complementation of light stress (HL)-induced SAA in the *rbohD* mutant with *RBOHD* driven by different tissue-specific promoters. Light stress‐induced systemic leaf cell injury (measured as electrolyte leakage) of wild type, *rbohD* and the different *rbohD*-complemented *Arabidopsis thaliana* plants is shown. Systemic leaves were either untreated and unstressed (Control) or subjected to a systemic light stress following a local pretreatment of a local leaf with light stress (SAA). Ten different plants each from two independent complemented lines for each construct were subjected to light stress and cell injury was determined by measuring electrolyte leakage from systemic leaves. Student t-test, SD, N=10, *p < 0.05. *Abbreviations used*: HL, high light; RBOHD, respiratory burst oxidase homolog D; CER, eceriferum; CAB, chlorophyll A/B binding protein; SCR, scarecrow; XCP, xylem cysteine peptidase; Sultr, sulfate transporter; EL, electrolyte leakage; SAA, systemic acquired acclimation. The experiments shown were conducted in parallel to the experiments shown in Figure 4 and are a repeat of the study reported previously (Zandalinas et al., 2020b), with similar results.

**Supplementary Figure S3.** Complementation of light (HL)- and heat (HS)-induced local and systemic ROS signaling in the *rbohD* mutant with *RBOHD* driven by the phloem- or mesophyll tissue-specific promoters, during stress combination. Representative images of whole-plant ROS levels in *rbohD* and *rbohD-*complemented *Arabidopsis thaliana* plants 20 min following a local light (HL)- or heat (HS)-treatments, or a combination of light- and heat-stress treatments applied to two leaves of the same plant (HL&HS; Zandalinas et al., 2020a; red circles) are shown. All experiments were repeated at least 3 times with similar results. Scale bar indicates 1 cm. *Abbreviations used*: HL, high light; HS, heat stress; RBOHD, respiratory burst oxidase homolog D; CAB, chlorophyll A/B binding protein; Sultr, sulfate transporter.

