## Supplemental Material for "Mesophyll cells mediate systemic reactive oxygen signaling during wounding or heat stress"

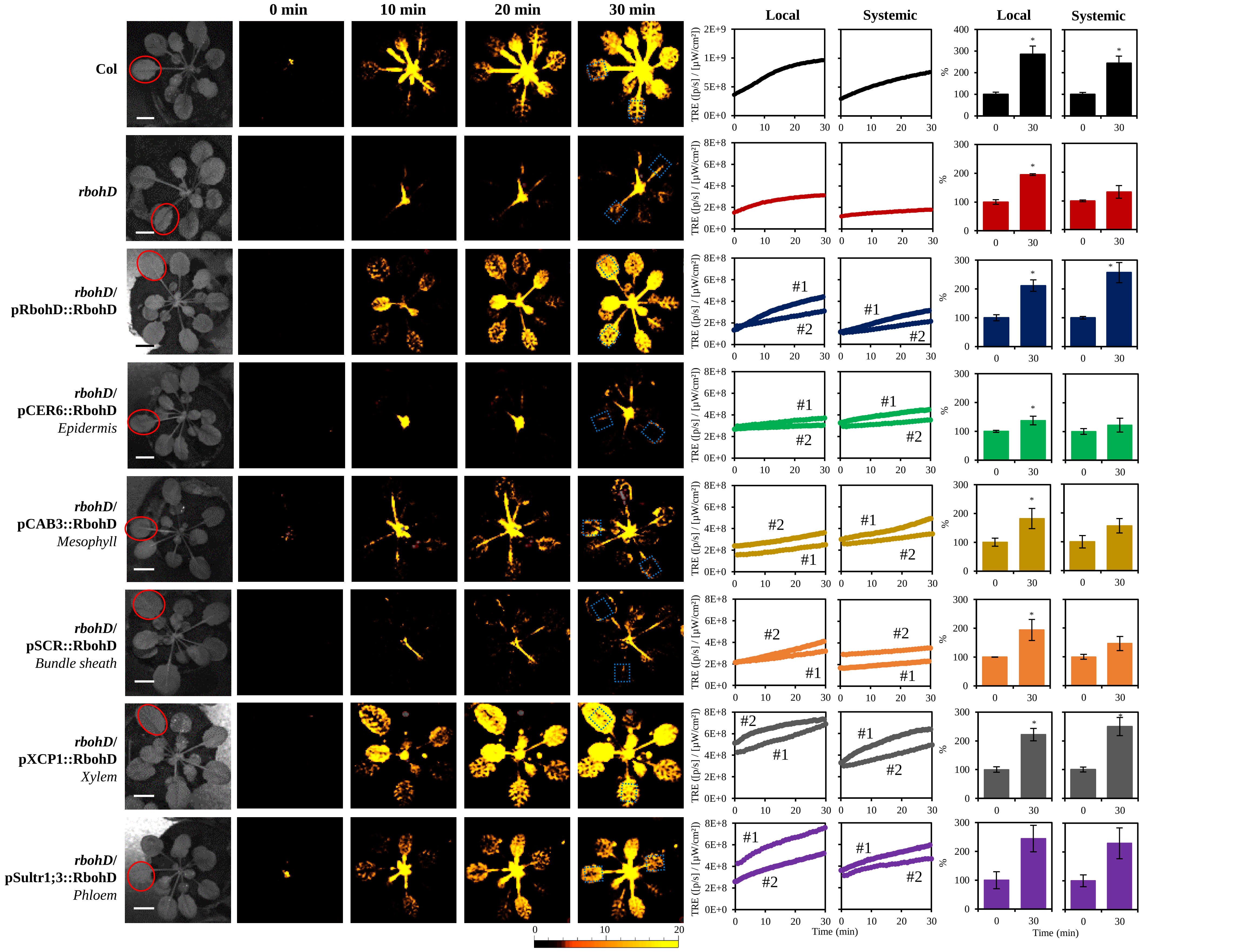


**Supplementary Figure S1.** Complementation of light (HL) stress-induced local and systemic ROS signaling in the *rbohD* mutant with *RBOHD* driven by different tissue-specific promoters. Representative time-lapse images of whole-plant ROS levels in wild type, *rbohD* and the different *rbohD* complemented *Arabidopsis thaliana* plants subjected to a local HL-stress treatment (red circles), are shown on left; representative line graphs showing continuous measurements of ROS levels in local and systemic leaves of wild type, *rbohD* and two independent homozygous complemented lines (#1 and #2), over the entire course of the experiment (0 to 30 min) are shown in the middle (ROIs for some of them are indicated with blue boxes); and statistical analysis of ROS levels in local and systemic leaves at 0 and 30 min is shown on right (Student t-test, SD, N=10, *$p < 0.05$). All experiments were repeated at least 3 times with similar results. Scale bar indicates 1 cm. *Abbreviations used*: HL, high light; RBOHD, respiratory burst oxidase homolog D; CER, eceriferum; CAB, chlorophyll A/B binding protein; SCR, scarecrow; XCP, xylem cysteine peptidase; ROI, region of interest; Sultr, sulfate transporter; TRE, total radiant efficiency. The experiments shown were conducted in parallel to the experiments shown in Figures 1 and 2 and are a repeat of the study reported previously (Zandalinas et al., 2020b), with similar results.

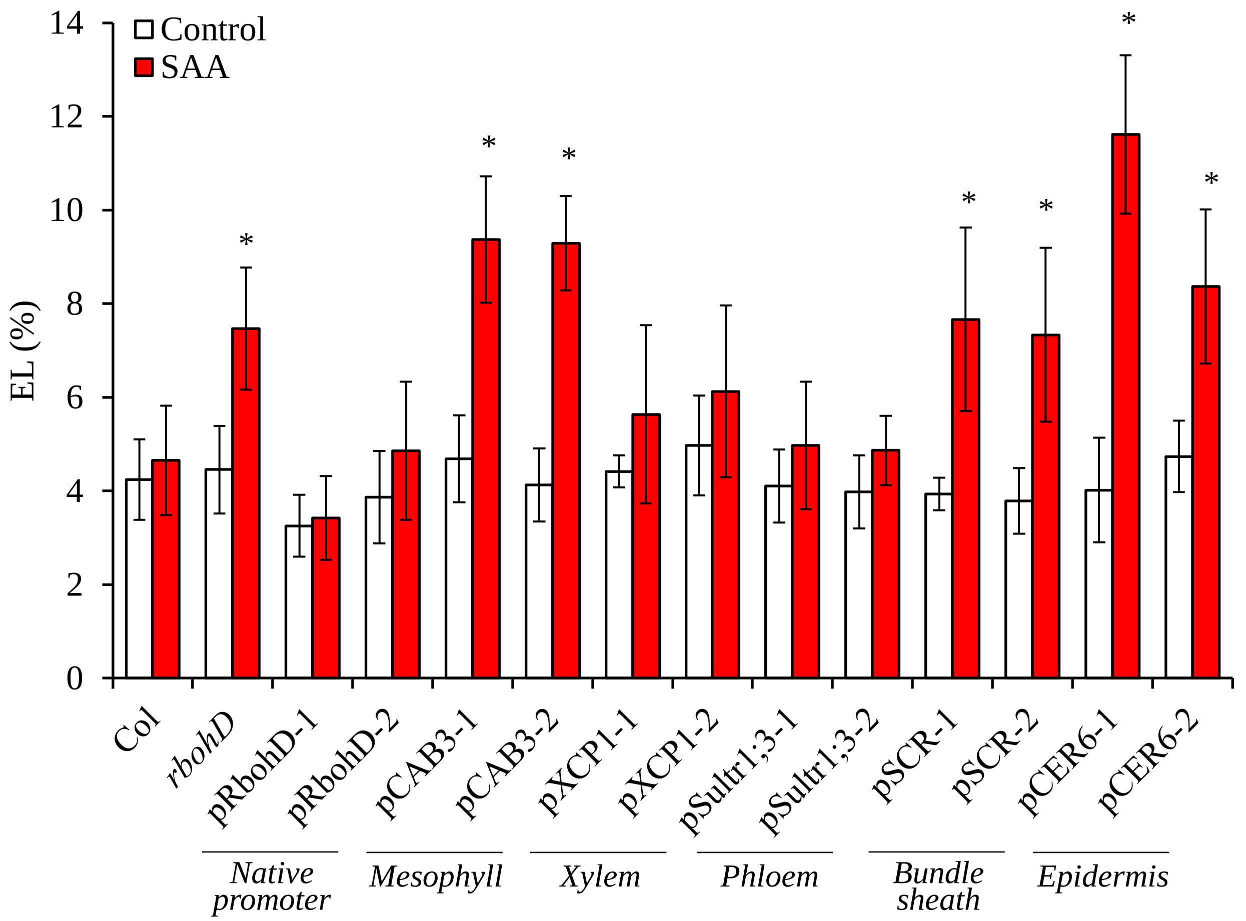


**Supplementary Figure S2.** Complementation of light stress (HL)-induced SAA in the *rbohD* mutant with *RBOHD* driven by different tissue-specific promoters. Light stress-induced systemic leaf cell injury (measured as electrolyte leakage) of wild type, *rbohD* and the different *rbohD*-complemented *Arabidopsis thaliana* plants is shown. Systemic leaves were either untreated and unstressed (Control) or subjected to a systemic light stress following a local pretreatment of a local leaf with light stress (SAA). Ten different plants each from two independent complemented lines for each construct were subjected to light stress and cell injury was determined by measuring electrolyte leakage from systemic leaves. Student t-test, SD, N=10, *$p < 0.05$. *Abbreviations used*: HL, high light; RBOHD, respiratory burst oxidase homolog D; CER, eceriferum; CAB, chlorophyll A/B binding protein; SCR, scarecrow; XCP, xylem cysteine peptidase; Sultr, sulfate transporter; EL, electrolyte leakage; SAA, systemic acquired acclimation. The experiments shown were conducted in parallel to the experiments shown in Figure 4 and are a repeat of the study reported previously (Zandalinas et al., 2020b), with similar results.

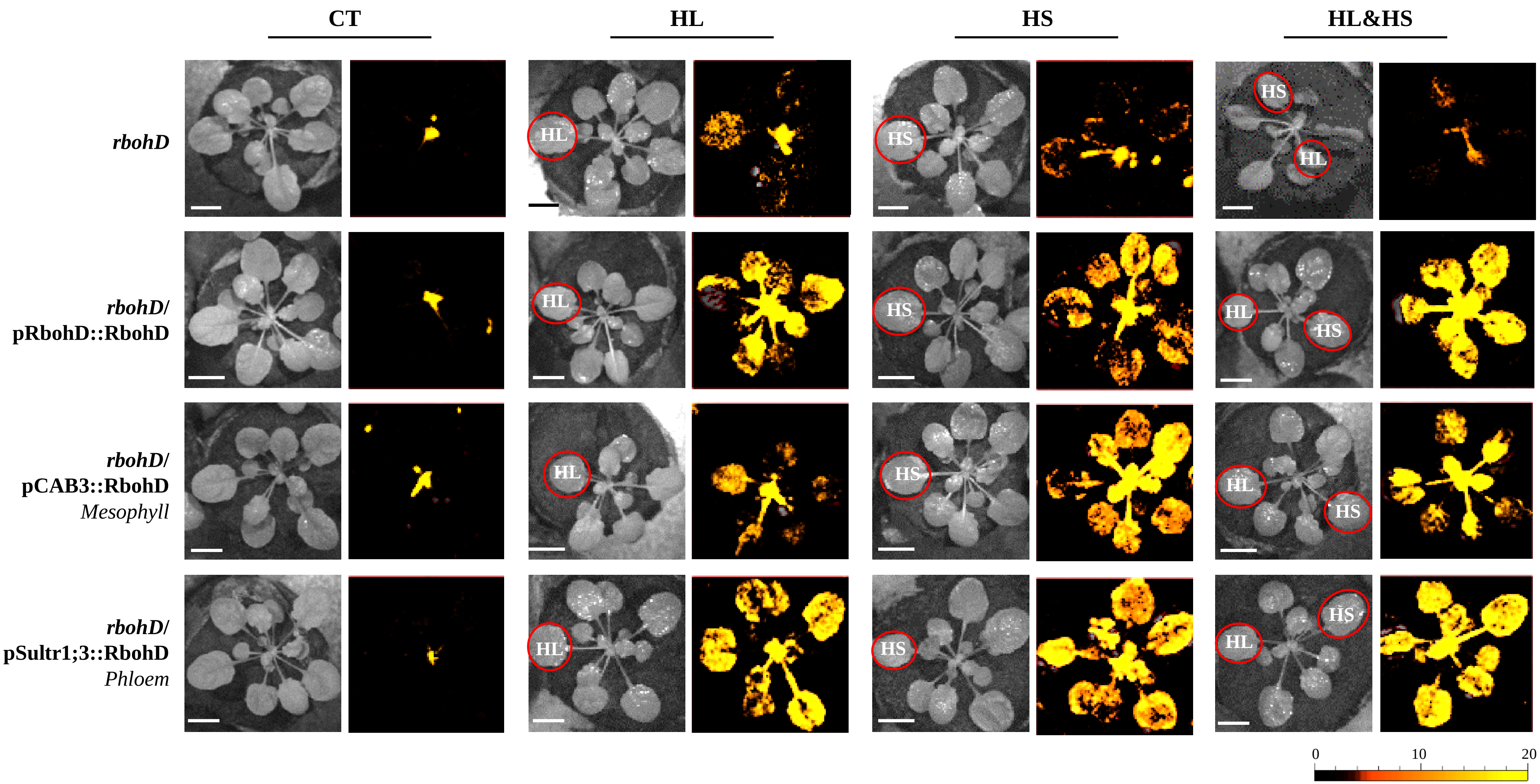


**Supplementary Figure S3.** Complementation of light (HL)- and heat (HS)-induced local and systemic ROS signaling in the *rbohD* mutant with *RBOHD* driven by the phloem- or mesophyll tissue-specific promoters, during stress combination. Representative images of whole-plant ROS levels in *rbohD* and *rbohD*-complemented *Arabidopsis thaliana* plants 20 min following a local light (HL)- or heat (HS)- treatments, or a combination of light- and heat-stress treatments applied to two leaves of the same plant (HL&HS; Zandalinas et al., 2020a; red circles) are shown. All experiments were repeated at least 3 times with similar results. Scale bar indicates 1 cm. *Abbreviations used*: HL, high light; HS, heat stress; RBOHD, respiratory burst oxidase homolog D; CAB, chlorophyll A/B binding protein; Sultr, sulfate transporter.

**Supplementary Table S1.** Transcript-specific primers used for relative expression analysis by RT-qPCR.

| Gene | Accession | Primer | |
|---|---|---|---|
| *JAZ5* | AT1G17380 | F<br>R | TCATCGTTATCCTCCCAAGC<br>CACCGTCTGATTTGATATGGG |
| *JAZ7* | AT2G34600 | F<br>R | GATCCTCCAACAATCCCAAA<br>TGGTAAGGGGAAGTTGCTTG |
| *Rap2.4* | AT1G78080 | F<br>R | TTTGCGCGGCTTAATTTCCC<br>TCACTGCGACGGTTGATGAA |
| *ERF2* | AT5G47220 | F<br>R | AACGAGCTGCGACTCAATGA<br>TCGACTTTAACCGCCGGAAA |
| *MYB30* | AT3G28910 | F<br>R | CTACCAATACTGGGCTGCTTAGAT<br>CTAGCTGAGGAAGTAGAGCGTCTT |
| *ZHD5* | AT1G75240 | F<br>R | CCACCAATCCAAGTCTCCCTC<br>GCTCGCCGCATGATTCTTTAG |
| *EF1α* | AT5G60390 | F<br>R | GAGCCCAAGTTTTTGAAGA<br>CTAACAGCGAAACGTCCCA |
